# Lipid-correlated alterations in the transcriptome are enriched in several specific pathways in the postmortem prefrontal cortex of Japanese patients with schizophrenia

**DOI:** 10.1101/2022.03.14.483348

**Authors:** Wataru Arihisa, Takeshi Kondo, Katsushi Yamaguchi, Junya Matsumoto, Hiroki Nakanishi, Yasuto Kunii, Hiroyasu Akatsu, Mizuki Hino, Yoshio Hashizume, Shumpei Sato, Shinji Sato, Shin-Ichi Niwa, Hirooki Yabe, Takehiko Sasaki, Shuji Shigenobu, Mitsutoshi Setou

**Author notes:** **Contact Information** Correspondence to: Mitsutoshi Setou, M.D., Ph.D., Director, International Mass Imaging Center, Hamamatsu University School of Medicine, 1-20-1 Handayama, Higashi-ku, Hamamatsu, Shizuoka 431-3192, Japan.

## Abstract

**Background:** Schizophrenia is a chronic relapsing psychiatric disorder that is characterized by many symptoms and has a high heritability. A previous study showed that specific lipid molecules belong to phosphatidylinositol (PI) and phosphatidylserine (PS) was reduced in the postmortem prefrontal cortex of patients with schizophrenia^1^. However, signaling pathways contributing to the lipid changes remain unknown. Here we performed two types of transcriptome analyses in patients with schizophrenia: an un-biased transcriptome analysis solely based on RNA-seq data and a correlation analysis between levels of gene expression and lipids.

**Methodology/Principal Findings:** RNA-Seq analysis was performed in the postmortem prefrontal cortex from 10 subjects with schizophrenia and 5 controls. Correlation analysis between the transcriptome and lipidome from 9 subjects which are the same samples in the previous lipidomics study^1^ (Table 1). Extraction of differentially expressed genes (DEGs) and further sequence and functional group analysis revealed changes of gene expression levels in phosphoinositide 3-kinase (PI3K)-Akt signaling and the complement system. In addition, a correlation analysis clarified alterations in several signaling/metabolic pathways including lipid-correlated genes, most of which are not found as DEGs in transcriptome analysis alone.

Table 1.
Characteristics of patients from whom postmortem brain samples were obtained. Abbreviations: PMI, postmortem interval, the time that has elapsed since a person has died; DOI, duration of illness, The samples used in correlation analysis are shown by black circles.

**Table 1. Characteristics of patients from whom postmortem brain samples were** 526 **obtained.**
| Sample | Diagnosis | Age (years) | Sex | PMI (hours) | DOI (years) | Correlation<br>Analysis |
| --- | --- | --- | --- | --- | --- | --- |
| S01 | SCZ | 87 | F | 5.0 | 49 |  |
| S02 | SCZ | 75 | M | 19.0 | 47 | • |
| S03 | SCZ | 68 | F | 16.0 | 40 | • |
| S04 | SCZ | 77 | F | 26.0 | 48 |  |
| S05 | SCZ | 66 | M | 6.0 | 36 |  |
| S06 | SCZ | 65 | F | 6.0 | 50 |  |
| S07 | SCZ | 70 | M | 31.0 | 39 | • |
| S08 | SCZ | 64 | M | 6.5 | 47 | • |
| S09 | SCZ | 71 | M | 17.5 | 48 | • |
| S10 | SCZ | 79 | F | 19.0 | 60 | • |
| **Average** |  | 72.2 |  | 15.2 | 46.4 |  |
| **SD** |  | 7.3 |  | 9.1 | 6.8 |  |
| C01 | CON | 80 | M | 2.0 |  |  |
| C02 | CON | 85 | F | 8.0 |  | • |
| C03 | CON | 89 | F | 4.0 |  | • |
| C04 | CON | 66 | M | 42.0 |  | • |
| C05 | CON | 76 | M | 6.0 |  |  |
| **Average** |  | 79.2 |  | 12.4 |  |  |
| **SD** |  | 8.9 |  | 16.7 |  |  |

**Conclusions:** This study provided results of the first integrated analysis of the schizophrenia-associated transcriptome and lipidome within the PFC and revealed that lipid-correlated alterations in the transcriptome are enriched in specific pathways including PI3K-Akt signaling.

## Introduction

Schizophrenia is a chronic relapsing psychiatric disorder that is characterized by many symptoms and affects an estimated one percent of the population globally. Hallucinations, delusions and abnormal behaviors are the most common symptoms of schizophrenia, but the symptoms vary from one person to another. Schizophrenia has high familial aggregation and is thus considered to arise from the interplay between genetic and environmental factors. Since the heritability is about 80%^2^, much research on the genomics of schizophrenia has been conducted worldwide. A genome-wide association study (GWAS) in many patients with schizophrenia revealed many risk genes, but the odds ratios were very low ^3^. Many candidate genes that explain the high heritability of schizophrenia were found in that GWAS, but that study also showed a complexity of the pathology of schizophrenia. The major functional gene categories suggested by that GWAS included neuronal, immune and histone pathways^4^.

Many studies have suggested that dysfunction of the prefrontal cortex (PFC) is associated with the pathology of schizophrenia, especially cognitive impairment^5–10^. Hirayama et al.^11^ used Japanese postmortem brain samples from Brodmann area 10 (BA10) in the PFC for comparing protein expression levels in patients with schizophrenia and those in controls. They showed that GNA13-ERK1-eIF4G2 signaling was downregulated and expression of CYFIP1 was decreased in patients with schizophrenia. CYFIP1 is involved in actin remodeling, and suppression of CYFIP1 expression impairs axon formation and synapse plasticity. Matsumoto et al.^1^ also used Japanese postmortem brain samples from BA10 in the PFC for quantitative and qualitative analyses of phospholipids in brain tissue of patients with schizophrenia. They found decreased levels of 16:0/20:4-phosphatidylinositol, PI(16:0/20:4), and 18:0/22:6-phosphatidylserine, PS(18:0/22:6), by using liquid chromatography electrospray ionization mass/mass spectrometry (LC-ESI/MS/MS) and imaging mass spectrometry (IMS). Another study showed abnormal distributions of phosphatidylserine (PS) and phosphatidylcholine (PC) in postmortem brain samples of patients with schizophrenia^12^. However, the factors that cause these lipid abnormalities and pathology of schizophrenia have not been elucidated.

It is assumed that an altered expression level of an mRNA in a disease is an indication for an abnormality in either the encoded protein or a biologically connected structure or function. In schizophrenia, transcriptomes have been most thoroughly studied with microarrays and a single microarray. Next-generation sequencing (NGS) has recently begun to replace microarray studies^13^. Deep sequencing approaches sequence the entire transcriptome multiple times and collect information on sequence variants and frequency of transcripts^14, 15^. Transcriptome analysis by NGS, so-called RNA-Seq, revealed that more than 2,000 genes displayed schizophrenia-associated alternative promoter usage and that more than 1,000 genes showed differential splicing (false discovery rate, FDR < 0.05)^16^. The first transcriptome study using many schizophrenia post-mortem brain samples pointed primarily to pathways related to synapse functions previously indicated by the results of a GWAS^17^.

In the present study, we analyzed the transcriptome in the PFC of patients with schizophrenia with consideration of the previous results of lipidome analysis into consideration. The PI3K-Akt signaling and the complement system were identified as pathways including DEGs from RNA-seq data alone. In addition, a correlation analysis between the transcriptome and lipidome revealed several interesting pathways containing lipid-correlated genes in the PFC of patients with schizophrenia.

## Results

### Identification of 119 differentially expressed genes (DEGs) in the PFC with schizophrenia

To prepare cDNA libraries, total RNA was extracted from post-mortem human brains from patients with schizophrenia and controls (Table 1). We used samples that were analyzed in a previous study^1^ to compare changes in gene expression and lipids. After a quality check, the cDNA libraries from patients with schizophrenia (N = 10) and controls (N = 5) were analyzed by using Illumina HiSeq1500 (see methods). The quality of FASTQ file was verified by the QRQC package in R ^18^.

The sequencing produced a total of 12,000 million bp fragments (sequence length range of 66 bp, 66 bp * each of the total sequences = 12,000 Mbp). These reads were mapped to the human genome using TopHat^19^. TopHat is a mapping tool that aligns sequencing reads to a reference genome for identifying splice junctions between exons. The average mapping rates was 98%, indicating a good quality of reads.

Analysis of differentially expressed genes (DEGs) by Cuffdiff^20^ revealed 119 significantly altered genes (FDR < 0.05) in schizophrenia samples compared to those in controls (Table S1). FDR is a widely used statistical method for multiple test correction. All gene expression values were calculated as FPKM (fragments per kilo base of exon per million mapped reads), a normalized measure for adjusting read counts by normalizing for gene length and total number of mapped reads for each sample. FPKM allow comparison at transcript levels across samples. Among the 119 DEGs identified, 88 genes were down-regulated and 32 genes were up-regulated. The fold change ranged from -2.39 to 4.57. RRAS, ZNF536, and CD14 were previous identified as genes located on 108 genetic loci in a GWAS^3^.

### Identification of schizophrenia-related pathways by enrichment analysis

To investigate the common features in the 119 DEGs, Go enrichment analysis was performed by DAVID^21, 22^. Five GO terms were significantly enriched at the stringent cut-off level of FDR < 0.05 (Table 2). These Go terms belong to Biological Process (BP) or Cellular Component (CC): negative regulation of endopeptidase activity (FDR = 0.038 < 0.05, gene count = 7), extracellular region (FDR = 0.0001, gene count = 27), extracellular matrix (FDR = 0.0003, gene count = 12), extracellular space (FDR = 0.0099, gene count = 21), and extracellular exosome (FDR = 0.03, gene count = 31). These results indicate that GO enrichment analysis is not sufficient to clarify the association of alterations in lipid levels in patients with schizophrenia.

**Table 2.**
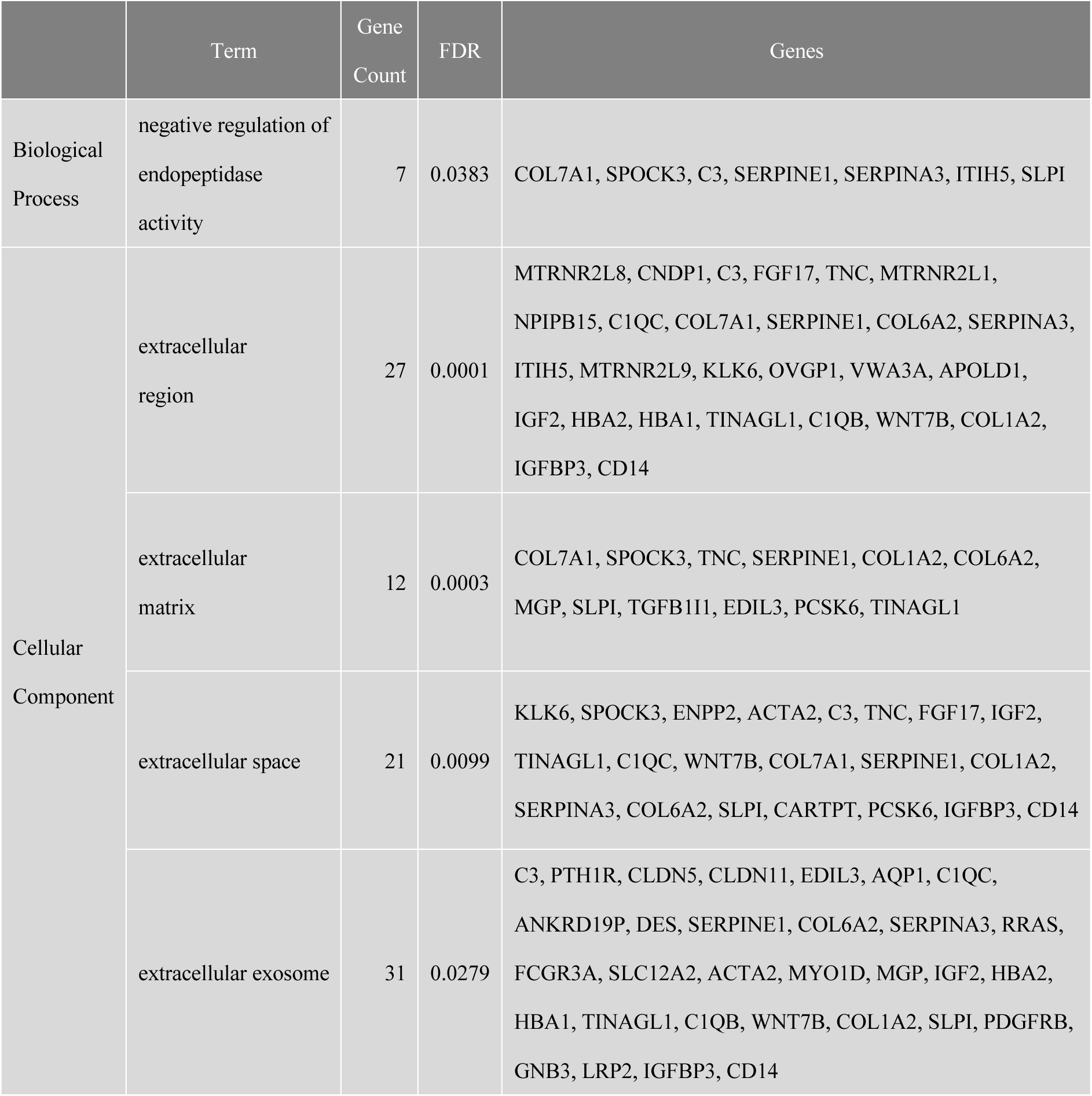
GO terms extracted from DEGs by an enrichment analysis tool (DAVID) Reported are the unique significantly enriched terms with FDR < 0.05 from 119 DEGs using the DAVID server.

To further explore the DEGs that belong to specific functional groups, we extracted signaling pathways to which the 119 DEGs belong by using KEGG pathway analysis^21, 22^ in the DAVID server (Table 3). KEGG (Kyoto Encyclopedia of Genes and Genomes) is the reference database for pathway mapping for associating genes and proteins with enzyme reactions and signaling pathways. We found seven significant (p-value < 0.05) pathways by DAVID^23^: Aminoacyl-tRNA biosynthesis (p-value = 0.0001, gene count = 6), Staphylococcus aureus infection (p-value = 0.0005, gene count = 5), Complement and coagulation cascades (p-value = 0.0113, gene count = 4), Systemic lupus erythematosus (p-value = 0.0127, gene count = 5), Pertussis (p-value = 0.0141, gene count = 4), PI3K-Akt signaling pathway (p-value = 0.0283, gene count = 7), and Chagas disease (p-value = 0.0333, gene count = 4). Gene count means the number of DEGs found in the pathway. These results of KEGG pathway analysis gave us a clue for finding the candidate pathways associated with lipid changes in schizophrenia.

**Table 3.** Pathways extracted from DEGs by an enrichment analysis tool (DAVID) Reported are the unique significantly enriched pathways with a p-value < 0.05 from 119 DEGs using the DAVID server.

| Term | Gene Count | p-value | FDR | Genes |
| --- | --- | --- | --- | --- |
| Aminoacyl-tRNA biosynthesis | 6 | 0.0001 | 0.01 | TRNE, TRNS1, TRNQ, TRNA, TRNY, TRNN |
| Staphylococcus aureus infection | 5 | 0.0005 | 0.03 | HLA-DQB1, C1QB, C3, FCGR3A, C1QC |
| Complement and coagulation cascades | 4 | 0.0113 | 0.32 | C1QB, C3, SERPINE1, C1QC |
| Systemic lupus erythematosus | 5 | 0.0127 | 0.32 | HLA-DQB1, C1QB, C3, FCGR3A, C1QC |
| Pertussis | 4 | 0.0141 | 0.32 | C1QB, C3, C1QC, CD14 |
| PI3K-Akt signaling pathway | 7 | 0.0283 | 0.54 | FGF17, TNC, COL1A2, COL6A2, PDGFRB, GNB3, DDIT4 |
| Chagas disease | 4 | 0.0333 | 0.54 | C1QB, C3, SERPINE1, C1QC |

The PI3K-Akt signaling pathway is one of the well-known schizophrenia-related pathways^24–26^. The activity of this signaling pathway is dependent on PI level because PIP_3_ is produced from PI via PIP_2_ ^27–29^. The other 5 pathways related to infection and immunity include complement genes (*C1QB, C1QC* and *C3*). The complement system is also thought to be involved in schizophrenia^30–32^. We therefore further analyzed gene expression changes in these pathways.

### Differences in expression levels of genes in schizophrenia for the PI3K-Akt signaling pathway

Seven DEGs, *FGF17, TNC, COL1A2, COL6A2, PDGFRB, GNB3* and *DDIT4*, were identified in the PI3K-Akt signaling pathway (Figure 1). All of the DEGs except for TNC were up-regulated in the brains of patients with schizophrenia. In the six up-regulated DEGs, error in the schizophrenia group varied widely compared with that in the control group. Gene expression levels were similar among samples in the control group but were different among samples in the schizophrenia group. A few patients with schizophrenia showed much higher expression levels those in the control group. In addition, the key genes of this pathway, *Akt1, Akt2* and *Akt3,* did not show significant differences. The DEGs in this pathway were limited to the upstream of this pathway. These results indicate that PI3K-Akt signaling may be associated with the pathology of schizophrenia, but the signaling did not seem to be critical in our samples.

**Figure 1.**
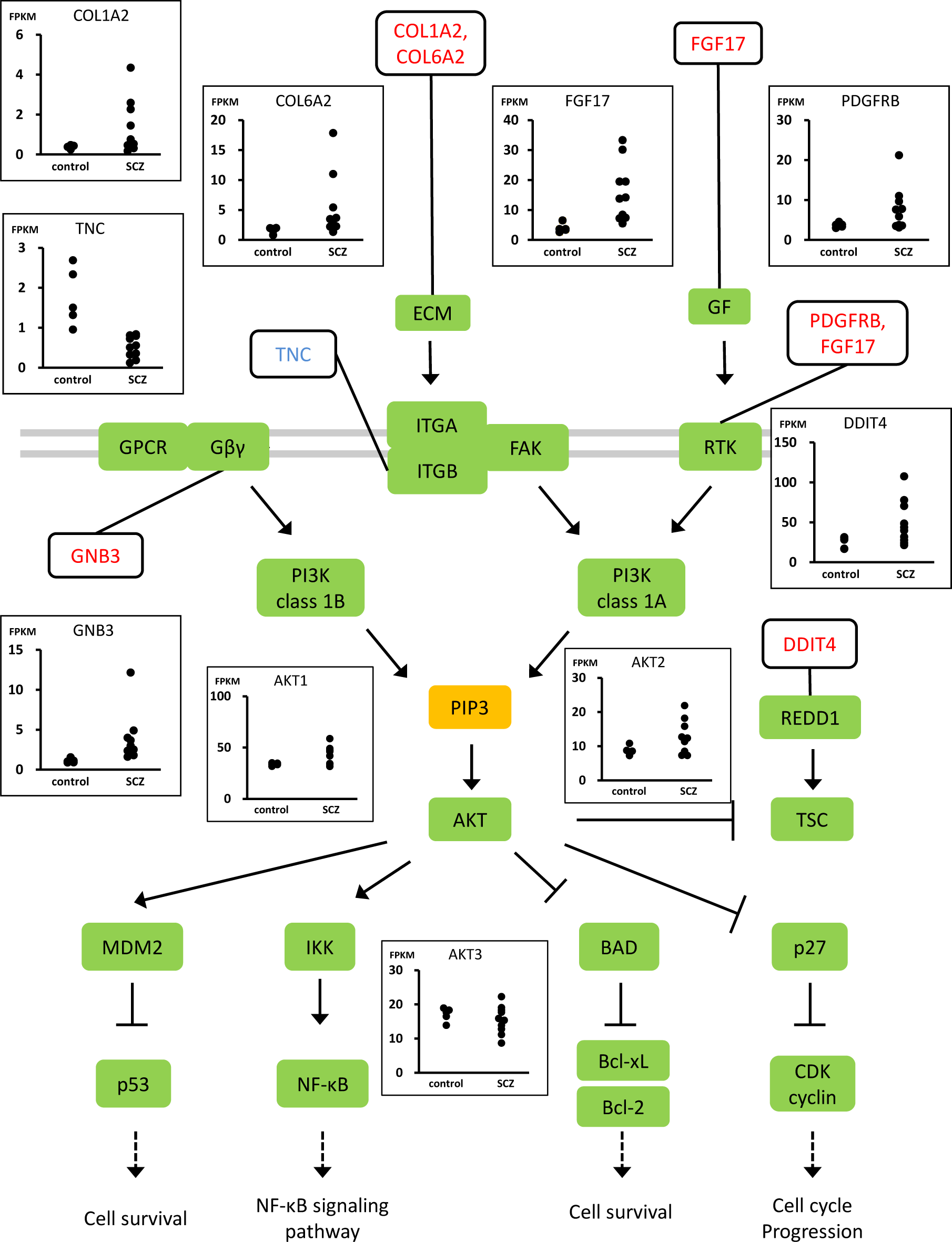
Expression levels of DEGs and non-DEGs in the PI3K-Akt signaling pathway. The KEGG Pathway for PI3K-Akt signaling with plotting of *FGF17, TNC, COL1A2, COL6A2, PDGFRB, GNB3, DDIT4, AKT1, AKT2,* and *AKT3*. The signal molecules involved in the PI3K-Akt pathway and proteins coded by DEGs found in this study are shown in rectangles. Expression levels of genes in each patient are presented as FPKM (vertical axis). Data points are grouped as schizophrenia patient or control (horizontal axis). The genes that were up-regulated are colored red and those that were down-regulated are colored blue. The green boxes show proteins and the orange box shows a lipid.

Differences in expression levels of genes in schizophrenia for the complement system Four DEGs, C1QB, C1QC, C3 and SERPINE1, were identified in the complement and coagulation cascades pathway (Figure 2). SERPINE1 was up-regulated and C3, C1QB and C1QC were down-regulated in the PFC of patients with schizophrenia. Other components belonging to the complement system such as C4 were not significantly different between the schizophrenia and control groups.

**Figure 2.**
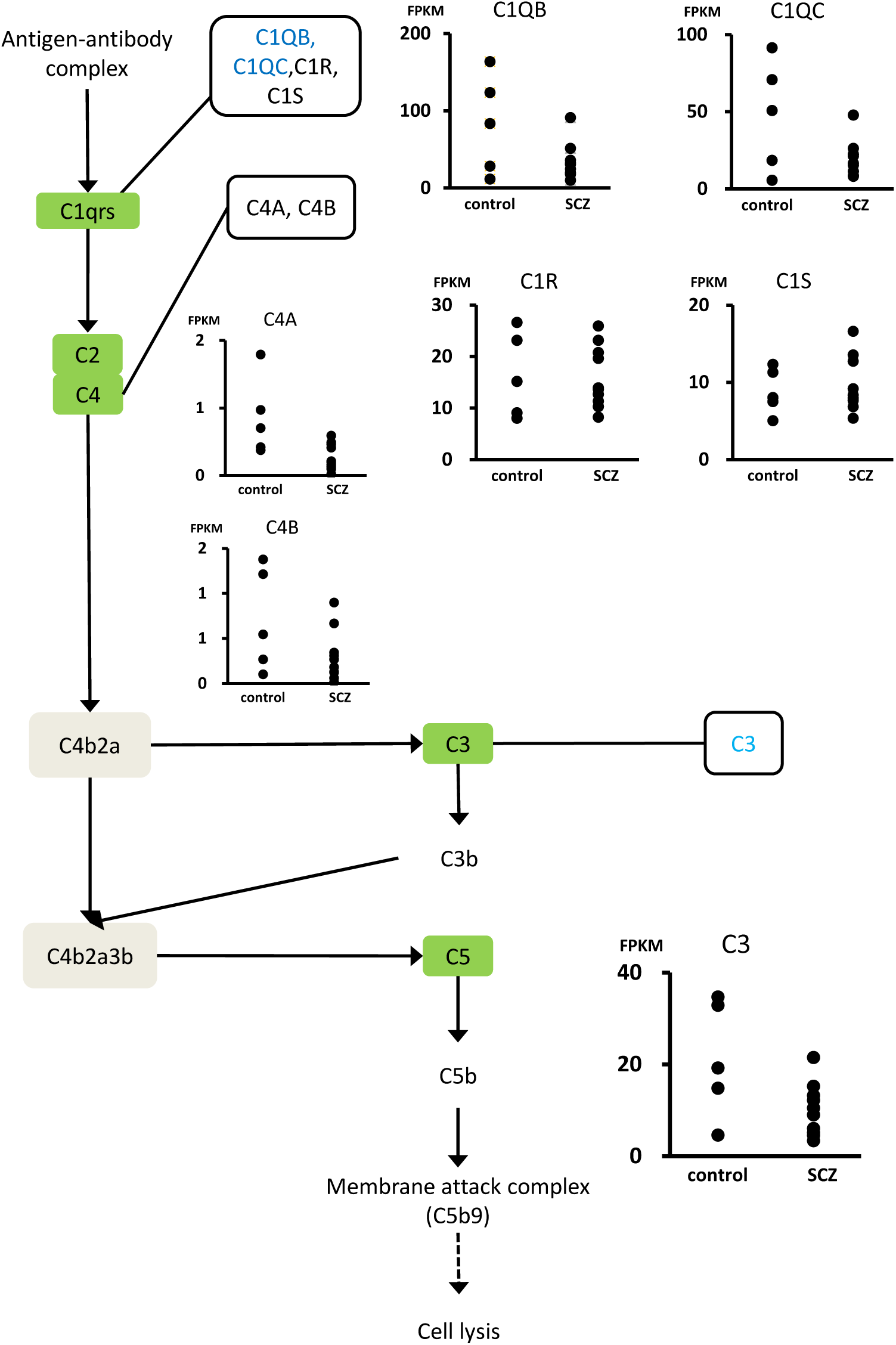
Expression levels of DEGs and non-DEGs in the classical pathway of complement cascade. The KEGG Pathway for classical pathway of complement cascade, with plotting *C1QB, C1QC, C1R, C1S, C3, C4A* and *C4B*. The signal molecules involved in the pathway and proteins coded by DEGs found in this study are shown in rectangles. Expression levels of genes in each patient are presented as FPKM (vertical axis). Data points are grouped as schizophrenia patient or control (horizontal axis). The genes that were up-regulated are colored red and those that were down-regulated are colored blue. The green boxes show proteins and the beige box shows convertase.

### Analysis of correlation of the transcriptome with changes of lipid levels in schizophrenia

Matsumoto et al. (2017)^1^ showed that PI(16:0/20:4) and PS(18:0/22:6) were significantly reduced in the same samples as those used in this study. Hence, we need to show the metabolic pathways around these lipids for investigating the related pathways of transcriptomics.

To explore correlated genes, we analyzed factors correlated with levels of PI(16:0/20:4) or PS(18:0/22:6) in the same samples. We selected 9 samples for which lipid data were available from the previous study^1^ (Table 1). Considering our sample size, we determined the correlation coefficients as ±0.8 (power = 0.8, significance level = 0.05) and extracted 389 genes with PI(16:0/20:4) and 623 genes with PS(18:0/22:6).

PI(16:0/20:4) and PS(18:0/22:6) shared most of correlated genes. To assess the pathways that these correlated genes belong to, we used DAVID again and found two pathways as correlated candidates with PI(16:0/20:4) (Table 4) and seven pathways as correlated candidates with PS(18:0/22:6) (Table 5) at a significance level of a p-value < 0.05. The two pathways with PI(16:0/20:4) were Ether lipid metabolism (p-value = 0.0009, gene count = 6) (Figure 3) and Cell adhesion molecules (CAMs) (p-value = 0.0104, gene count = 8) (Figure 4). The seven pathways with PS(18:0/22:6) were Ether lipid metabolism (p-value = 0.0005, gene count = 8) (Figure 3), Sphingolipid metabolism (p-value = 0.0035, gene count = 7), Lysosome (p-value = 0.005, gene count = 11), Axon guidance (p-value = 0.0197, gene count = 10), Endocytosis (p-value = 0.0435, gene count = 14), PI3K-Akt signaling pathway (p-value = 0.0478, gene count = 18) (Figure 5) and Cell adhesion molecules (CAMs) (p-value = 0.037, gene count = 10) (Figure 4). Ether lipid metabolism and Cell adhesion molecules were found as common pathways for PI(16:0/20:4) and PS(18:0/22:6).

**Figure 3.**
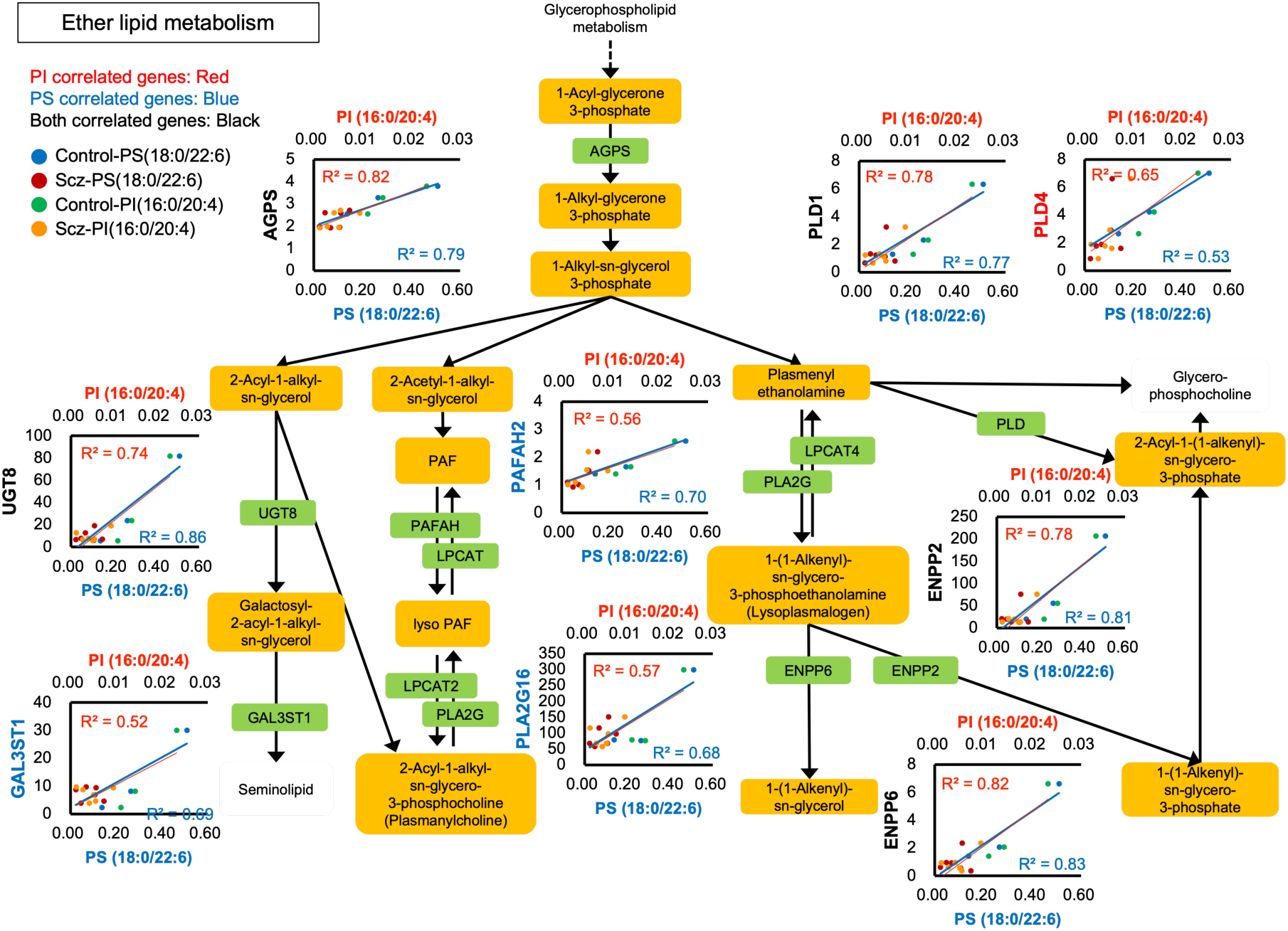
Correlations of gene expressions and lipid levels in Ether lipid metabolism. The KEGG Pathway for ether lipid metabolism with scatterplots of genes and lipids. The signal molecules involved in the pathway and proteins found in this study are shown in rectangles. The genes correlated to PI(16:0/20:4), PS(18:0/22:6) or both of them are shown in red, blue or black, respectively. The vertical axis of the scatterplot indicates expression levels of genes in each patient as FPKM. The upper label of horizontal axis indicates PI(16:0/20:4) levels and the lower label of horizontal axis indicates PS(18:0/22:6) levels. These values were calculated as relative intensities to those of internal standard (PC 14:0/14:0). Data points are classified as following 4 groups: blue, control group in PS(18:0/22:6); red, schizophrenia group in PS(18:0/22:6); green, control group in PI(16:0/20:4); orange, schizophrenia group in PI(16:0/20:4). The regression lines for PI(16:0/20:4) or PS(18:0/22:6) are shown in red or blue, respectively. Some nodes in the pathway are not shown due to space limitations.

**Figure 4.**
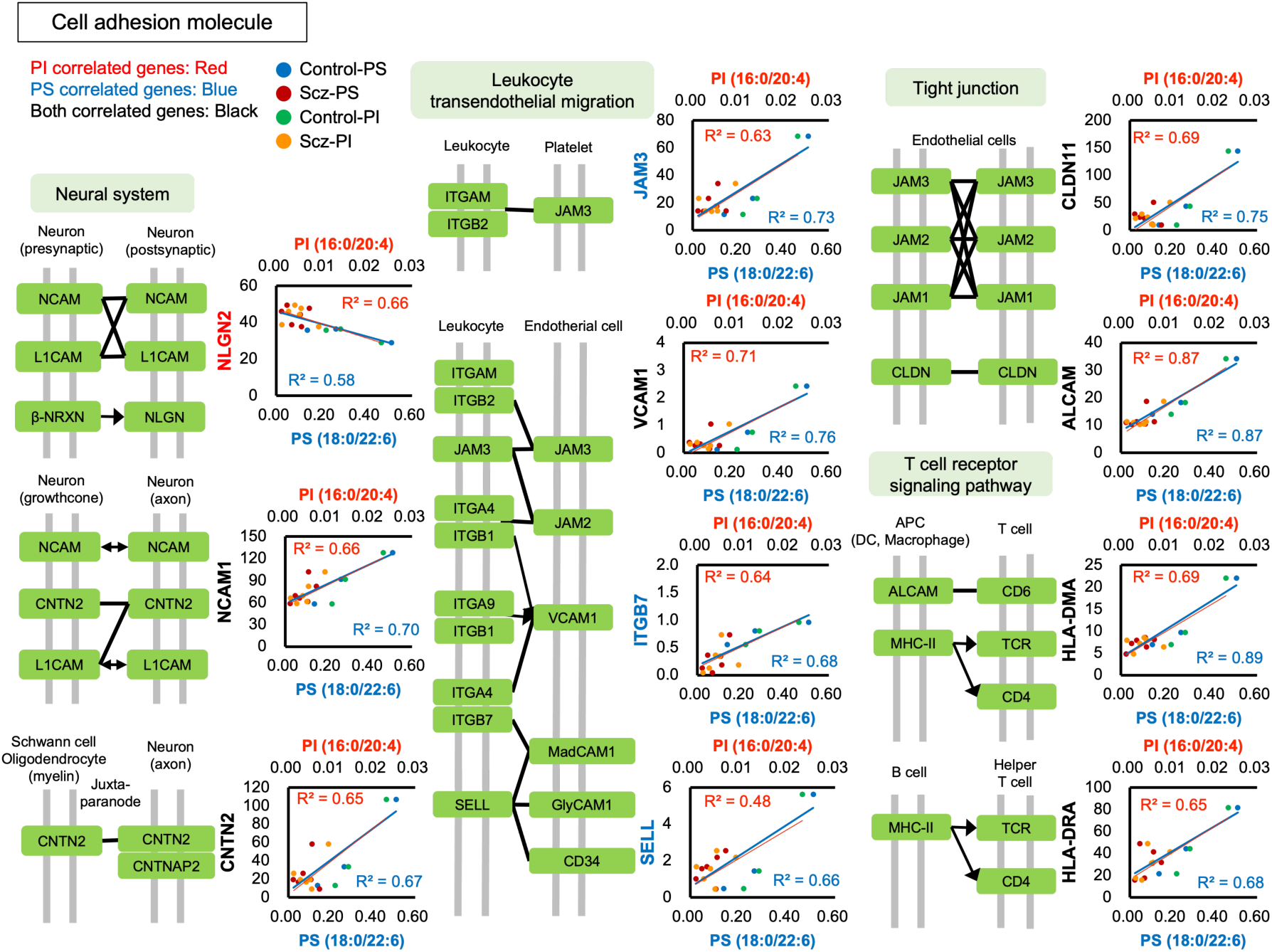
Correlations of gene expressions and lipid levels in Cell adhesion molecules. The KEGG Pathway for Cell adhesion molecules with scatterplots of genes and lipids. Black bars indicate protein-protein interactions on the surfaces of indicated cell types. The scatterplots are described same as in Figure 3.

**Figure 5.**
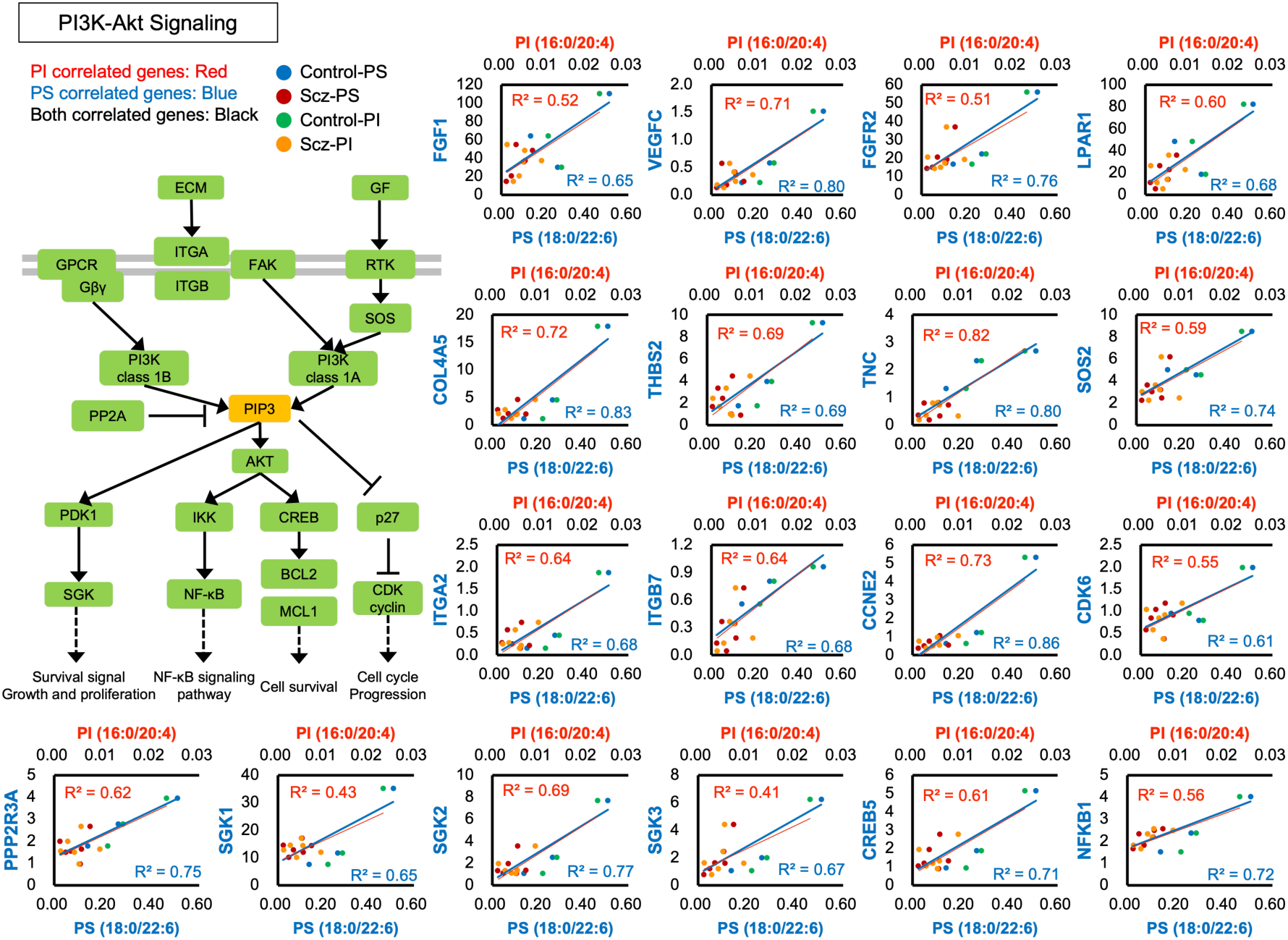
Correlations of gene expressions and lipid levels in PI3K-Akt signaling pathway. The KEGG Pathway for PI3K-Akt signaling pathway with scatterplots of genes and lipids. The molecules and scatterplots are described same as in Figure 3.

**Table 4.** Pathways extracted from genes correlated with PI(16:0/20:4) by an enrichment analysis tool (DAVID) Reported are the unique significantly enriched pathways with a p-value < 0.05 from 389 genes correlated with PI(16:0/20:4) using the DAVID server. Genes shown in red indicate inverse correlations with PI(16:0/20:4). DEGs identified in the RNA-seq analysis are shown in bold type.

| Term | Gene Count | p-value | FDR | Genes |
| --- | --- | --- | --- | --- |
| Ether lipid metabolism | 6 | 0.0009 | 0.15 | **UGT8**, AGPS, **ENPP2**, PLD4, ENPP6, PLD1 |
| Cell adhesion molecules (CAMs) | 8 | 0.0104 | 0.86 | **CLDN11**, HLA-DMA, **NLGN2**, VCAM1, ALCAM, HLA-DRA, CNTN2, NCAM1 |

**Table 5.**
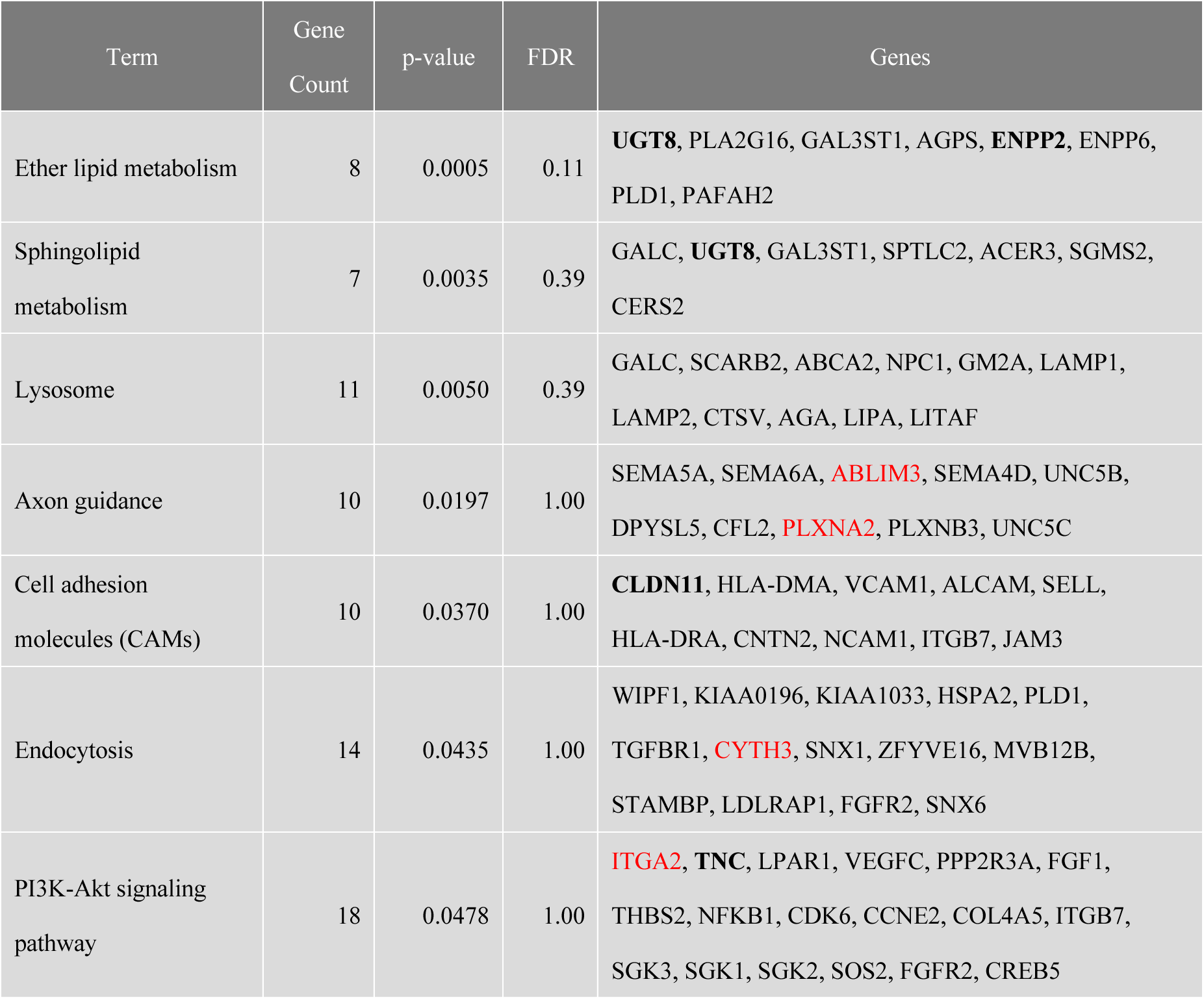
Pathways extracted from genes correlated with PS(18:0/22:6) by an enrichment analysis tool (DAVID) Reported are the unique significantly enriched pathways with a p-value < 0.05 from 623 genes correlated with PS(18:0/22:6) using the DAVID server. Genes shown in red indicate inverse correlations with PS(18:0/22:6). DEGs identified in the RNA-seq analysis are shown in bold type.

| Term | Gene Count | p-value | FDR | Genes |
| --- | --- | --- | --- | --- |
| Ether lipid metabolism | 8 | 0.0005 | 0.11 | **UGT8**, PLA2G16, GAL3ST1, AGPS, **ENPP2**, ENPP6, PLD1, PAFAH2 |
| Sphingolipid metabolism | 7 | 0.0035 | 0.39 | GALC, **UGT8**, GAL3ST1, SPTLC2, ACER3, SGMS2, CERS2 |
| Lysosome | 11 | 0.0050 | 0.39 | GALC, SCARB2, ABCA2, NPC1, GM2A, LAMP1, LAMP2, CTSV, AGA, LIPA, LITAF |
| Axon guidance | 10 | 0.0197 | 1.00 | SEMA5A, SEMA6A, **ABLIM3**, SEMA4D, UNC5B, DPYSL5, CFL2, **PLXNA2**, PLXNB3, UNC5C |
| Cell adhesion molecules (CAMs) | 10 | 0.0370 | 1.00 | **CLDN11**, HLA-DMA, VCAM1, ALCAM, SELL, HLA-DRA, CNTN2, NCAM1, ITGB7, JAM3 |
| Endocytosis | 14 | 0.0435 | 1.00 | WIPF1, KIAA0196, KIAA1033, HSPA2, PLD1, TGFBR1, **CYTH3**, SNX1, ZFYVE16, MVB12B, STAMBP, LDLRAP1, FGFR2, SNX6 |
| PI3K-Akt signaling pathway | 18 | 0.0478 | 1.00 | **ITGA2**, **TNC**, LPAR1, VEGFC, PPP2R3A, FGF1, THBS2, NFKB1, CDK6, CCNE2, COL4A5, ITGB7, SGK3, SGK1, SGK2, SOS2, FGFR2, CREB5 |

## Discussion

In this study, we analyzed the transcriptome in the PFC of patients with schizophrenia for finding the cause of changes in PI(16:0/20:4) and PS(18:0/22:6)^1^. Our pathway analysis based on RNA-seq data revealed DEGs related to PI3K-Akt signaling and the complement system in the PFC of patients with schizophrenia. Furthermore, correlation analyses between the transcriptome and lipidome identified several candidate pathways that were altered in schizophrenia.

PI3K-Akt signaling regulates cell survival, proliferation and apoptosis as well as synaptic formation and plasticity. Many previous studies have indicated a relationship between the PI3K-Akt signaling pathway and schizophrenia^33–36^. Enriquez-Barreto et al. proposed that such a disruption of this pathway is involved in the pathology of autism and schizophrenia^33^. Involvement of the disruption of the pathway was also suggested by other studies on schizophrenia^37, 38^. Although our sample size was very small, we can get this common pathway from both transcriptome analysis alone and correlation analysis with lipidome (as discussed below), further supporting the hypothesis that the PI3K-Akt signaling pathway plays an important role in the pathology of schizophrenia.

The expression levels of DEGs in the PI3K-Akt signaling pathway varied among the samples (Figure 1), reflecting the aspect of individuality in the pathology of schizophrenia. Notably, the expression levels of DEGs related to the PI3K-Akt signaling pathway were increased in brain samples with low levels of PI^1^, which are controversial to decreased expression level of PI3K and Akt and activation of the PI3K-Akt signaling pathway in schizophrenia^33^. These increased levels of DEGs do not seem to contribute to the PI(16:0/20:4) and PS(18:0/22:6) reductions because these DEGs are not directly involved in biosynthesis of these lipids. It is thought that these DEGs work in substitution for rather than the cause of the low activity of the PI3K-Akt signaling pathway. On the other hand, the expression of TNC was dramatically decreased in all patients. TNC is a gene that codes for tenascins, which are extracellular matrix glycoproteins, and is involved in neuroplasticity, and it has shown to be related to Alzheimer disease^39^. The decreased level of TNC in brains with a low PI level may exacerbate the dysfunction of the PI3K-Akt signaling pathway. To understand the role of this pathway in schizophrenia, further experiments such as experiments using a mouse model in which the pathway is suppressed are needed^40^.

A possible association between schizophrenia and the immune system has been indicated by numerous studies has been discussed as a potential mechanism for schizophrenia ^30, 41^. The complement system has recently emerged as genes located in MHC region which has strong association in schizophrenia^31, 32, 42–47^. Our results showed significant reductions of *C3, C1QA* and *C1QB* in schizophrenia (Figure 2), in contrast to the up-regulation of complements observed in previous studies. The reason for these controversial observations is currently unknown. However, some studies also showed reduced levels of serum complement proteins in schizophrenia^48–50^. Considering the results of two studies on Asian patients with schizophrenia, racial/ethnic factors may explain the different contributions of the complement system to the pathology of schizophrenia. A previous study also revealed up-regulation of several genes in the category belonging to the complement system including *C1R, C1S, C7, FCN3, SERPING1, C4A* and *CFI*^30^. However, in our samples, the expression levels of these genes were not different between the schizophrenia and control groups.

Our pathway analysis based on the genes correlated with lipid levels further demonstrated additional pathways that are possibly altered in schizophrenia, most of which were not identified by transcriptome analysis alone. PI(16:0/20:4) and PS(18:0/22:6) shared correlated genes for lipid ether metabolism (Figure 3) and cell adhesion molecules (CAMs) (Figure 4). The reduced levels of genes for lipid ether metabolism may contribute to the reduction of PI(16:0/20:4) and PS(18:0/22:6) because *PLDs* and *ENPP2* (a gene coding for autotaxin) regulate phosphatidic acid (PA) metabolism, which is a common source of phospholipid synthesis^51–53^. The class- and acyl chain-specific reduction in schizophrenia^1^ indicates additional factors for changes in PI(16:0/20:4) and PS(18:0/22:6). In addition, PA is known to be a regulator for mTOR^54^, implying a possible link to defects in the PI3K-Akt pathway^55^. Further assessments of PA would be helpful to understand these observations and pathologies in schizophrenia. Indisputably, CAMs are critical for the development and functions of the brain, including the plasticity of synaptic connections^56^. To date, to our knowledge, there has been no report showing that CAMs directly regulate phospholipid synthesis. Thus, defects in CAMs may reflect the lipid-related pathologies in this disorder rather than the machinery of the lipid changes. Consistent with our data, recent human genetic studies have suggested the involvement of CAMs in neuropsychiatric disorders including schizophrenia^57^, implying that loss of CAMs is one of the general features in the pathology of schizophrenia. A recent study using *CLDN11*-null mice showed that *CLDN11* (identified as one of the DEGs in this study) plays a role in the regulation of neurotransmission and behavior^58^.

PS(18:0/22:6) has additional correlated pathways: Sphingolipid metabolism, Lysosome, Axon guidance, Endocytosis and PI3K-Akt signaling pathway. Notably, PS(18:0/22:6), but not PI(16:0/20:4), has correlated genes belonging to the PI3K-Akt pathway (Figure 5). This observation is consistent with the results of a study suggesting that PS contributes to activation of the PI3K-Akt signaling^59^. Importantly, decreased levels of docosahexanoic acid (DHA)-containing PS seem to be associated with cognitive dysfunctions^60, 61^. In addition, our data indicate the possibility that reduction of TNC (found as one of the DEGs in this study) also contributes to dysregulation of PI3K-Akt signaling by PS(18:0/22:6) reduction. In contrast, DEGs related to the complement system were not extracted as correlated genes, indicating that the reductions of *C1QA, C1QB* and *C3* are independent of changes in PI(16:0/20:4) and PS(18:0/22:6). The other PS(18:0/22:6)-specific correlated genes in the sphingolipid metabolism, lysosome, endocytosis and axon guidance groups do not include DEGs, indicating ambiguous changes in schizophrenia. However, these pathways may have a modest disturbance because most of these genes showed reduced levels synchronously in schizophrenia. These pathways are known to be important for functions of the brain and their associations with schizophrenia have been discussed^62–69^. Further study is needed to clarify the mechanisms for these changes in levels of gene expression that are correlated with lipid alterations.

Taken together, we succeeded in identifying the pathway and DEGs that are related to the pathology of schizophrenia by combining the previous study on lipids. Schizophrenia is considered to be a syndrome, so it is difficult to investigate the pathology of schizophrenia by only single omics analysis. Indeed, in this study, we could not find a single factor to explain schizophrenia by transcriptome analysis. In addition, the causal relationship between the transcriptome and lipidome remains unknown because the data from a postmortem brain are single point data. However, intersection between transcriptome and lipidome successfully extracted several reasonable pathways that seem to be altered in the brains of patients with schizophrenia. Most of the lipid-correlated genes in these pathways were not identified as DEGs in transcriptome analysis alone, indicating that the combination of multiple omics study could be a helpful method for understanding complex disorders such as schizophrenia. Further studies on these identified pathways should provide useful insights into the pathology of schizophrenia.

## Materials & Methods

### Human brain tissue samples

Post-mortem brain samples from BA10 in the PFC of patients who had been diagnosed with schizophrenia were obtained from the Post-mortem Brain Bank of Fukushima for Psychiatric Research (Fukushima, Japan). Control samples were obtained from Choju Medical Institute, Fukushimura Hospital. This study, including the use of post-mortem human brain tissue, was approved by the Ethics Committee of Fukushima Medical University, Fukushimura Hospital and Hamamatsu University School of Medicine and complied with the Declaration of Helsinki. All procedures were carried out with the informed written consent of the next of kin. All patients diagnosed with schizophrenia had fulfilled the diagnostic criteria established by the American Psychiatric Association (Diagnostic and Statistical Manual of Mental Disorders: DSM-IV).

### Library preparation and sequencing

Two hundred fifty ng of total RNA from each of the brain samples was used for library preparation with the Illumina TruSeq RNA kit (Illumina, USA) by a half-scale method of the manufacturer’s instructions. The obtained libraries were evaluated using the KAPA library quantification kit (KAPA Biosystems, USA) and Bioanalyzer High Sensitivity (Agilent Technologies, USA). One sample from the control group was excluded from sequencing due to its low quality. Finally, the libraries of multiplexes (control, 5 samples; schizophrenia, 10 samples) were pooled and analyzed using Illumina HiSeq1500 (Illumina).

### Pre-processing and mapping of RNA-Seq reads using TopHat2

We extracted 66-bp length reads using Illumina HiSeq 1500. The reads assessed by Phred quality score were used to generate FASTQ files. These FASTQ files were then mapped to NCBI H. sapiens reference genome (GRCH38) using TopHat2 v2.11 with default parameters^19^, which calls Bowtie2 v2.2.5. NCBI H. sapiens GRCH38 was downloaded from iGenome (http://jp.support.illumina.com/sequencing/sequencing_software/igenome.html) and used as the reference genome. TopHat2 is a splice-aware aligner for splicing junctions that splits reads into short segments, and then the short segments of reads are mapped to the reference genome by Bowtie2. After initial mapping, TopHat2 splits the unmapped reads and aligns those reads again with possible splices.

### Transcript quantification using Cufflinks

Transcript quantification was accomplished by using Cufflinks v2.2.1, which includes Cuffdiff and measures the relative abundance of transcripts. Gene expression levels were quantified as FPKM.

## Acknowledgements

We thank the members of Advanced Research Facilities and Services (HUSM) and M. Matsumoto (National Institutes of Natural Sciences). We also thank Y. Sugiyama and S. Takano for secretarial assistance and the members of AMED-CREST “the optolipidomics project” for valuable discussion. We also thank SES translation and proofreading service (Sapporo, Japan) for English language editing. This study was supported by a KAKENHI Grant-in-Aid for JSPS Fellows (Grant Number 26-5334 to T.K.), Grant-in-Aid for Early-Career Scientists (Grant Number JP18K15063 to T.K.), Grants-in-Aid (Grant Number JP15H05899, Grant Number JP17H03980, Grant Number JP20H03433 to T.S.) and AdAMS (Grant Number JP16H06276) from the Japan Society for the Promotion of Science (JSPS), a CREST grant (Grant Number 921910520 to M.S., Grant Number JP18gm0910004 to T.K. and Grant Number JP18gm0710002h0006 to T.S.) and the Strategic Research Program for Brain Sciences (Grant Number JP21wm0425019 to H.Y. and Grant Number JP21dm020707 to Y.K.) from the Japan Agency for Medical Research and Development (AMED). This work was also supported by Ministry of Education, Culture, Sports, Science, and Technology of Japan (MEXT) for promoting public utilization of advanced research infrastructure (Imaging Platform, Grant Number JPMXS0410300221) and Grant-in-Aid for Scientific Research on Innovative Areas (Grant Number JP21H00180 to Y.K.).

## Author contributions

W.A., T.K., K.Y., and S. Shigenobu contributed to RNA-seq. J.M., Y.K., H.A., Y.H., M.H., S.N., and H.Y. contributed to the sample preparation of post-mortem human brains. H.N. and T.S. contributed to LC-MS. W.A., T.K., J.M., and Shumpei Sato performed the data analysis. W.A. and T.K. prepared a draft of the manuscript, which was validated by all of the other authors. M.S. and Shinji Sato supervised the entire project.

## Competing Financial Interests

There is no competing financial interest.

## Supplemental Figures and legends

**Table S1.**
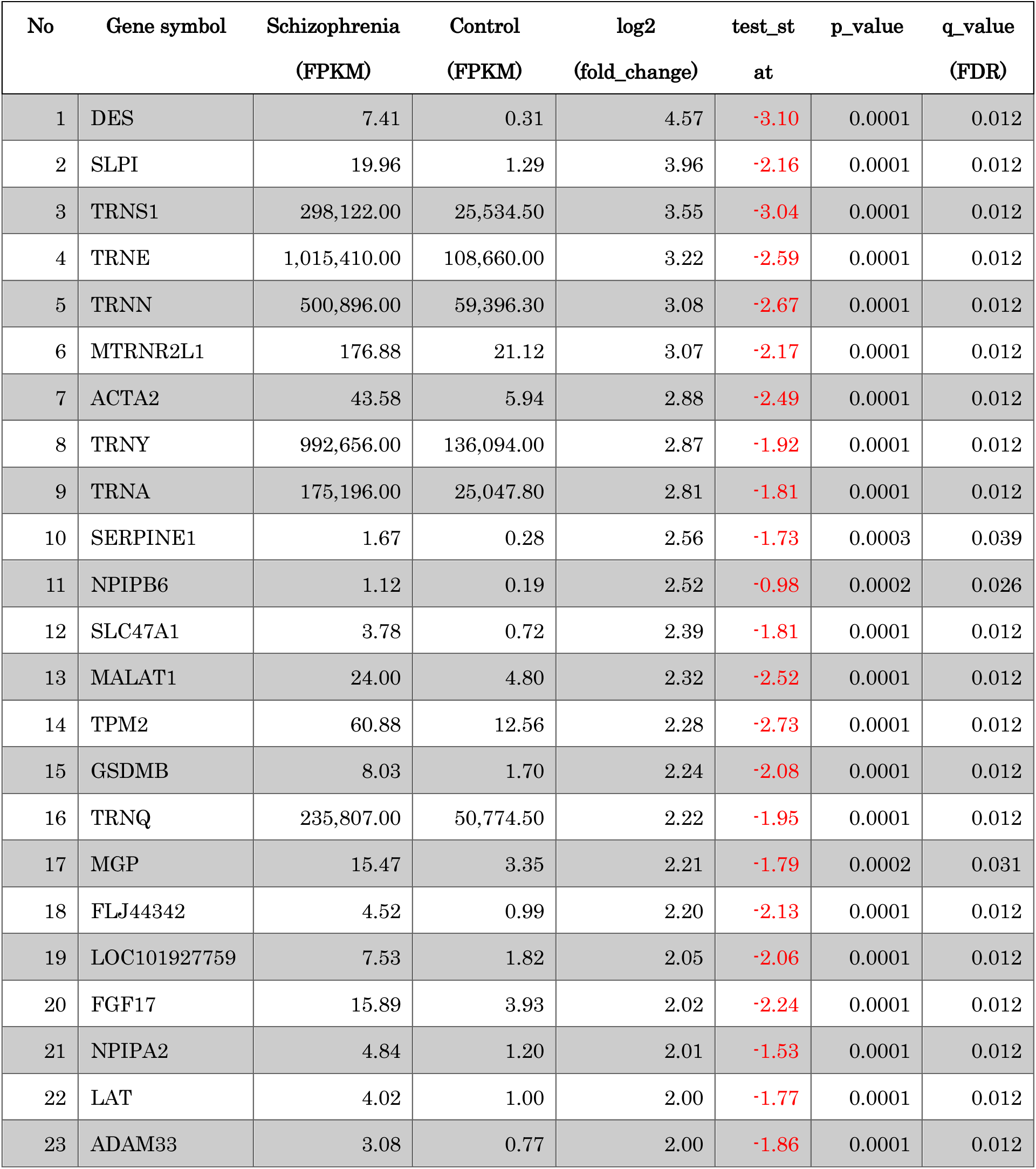
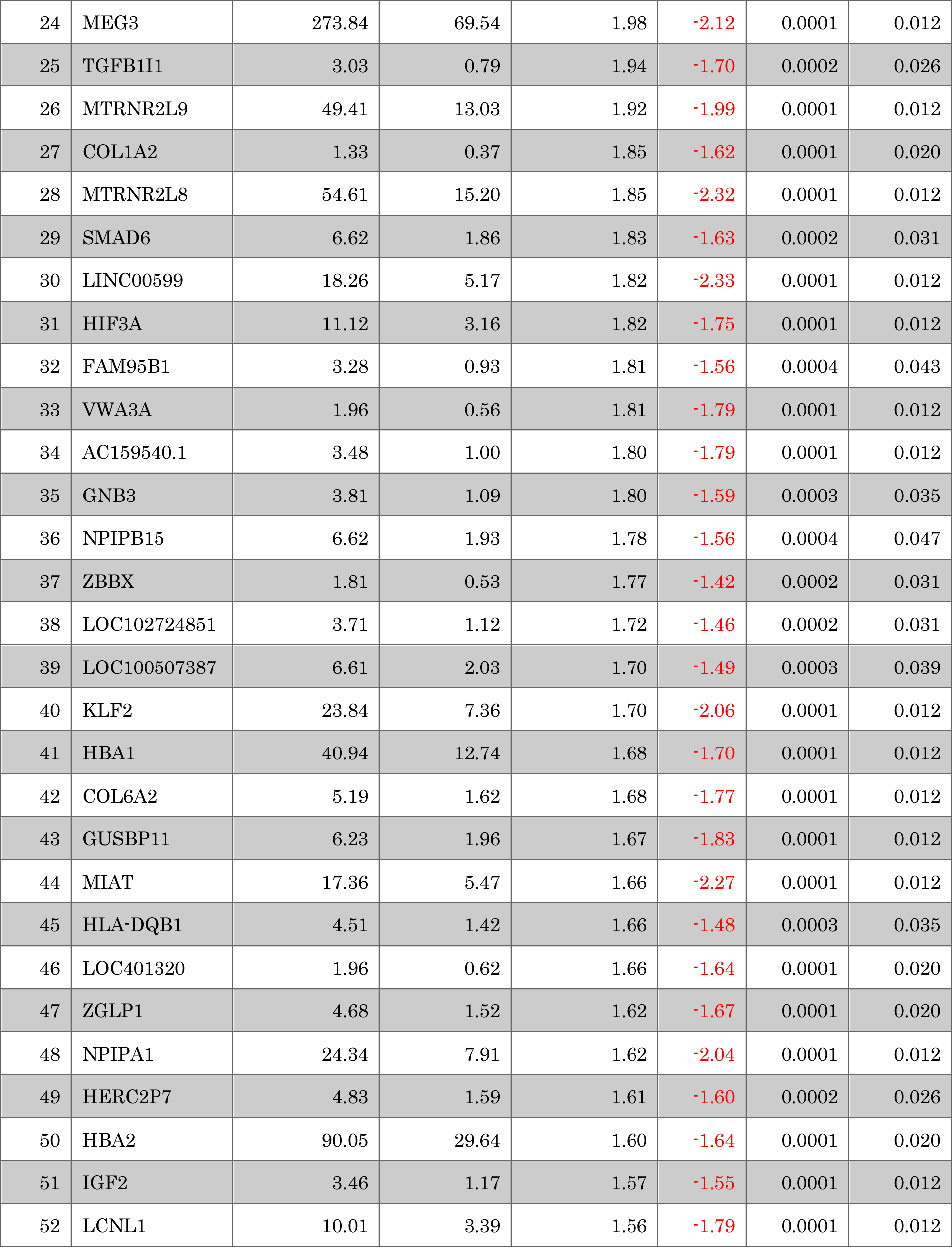
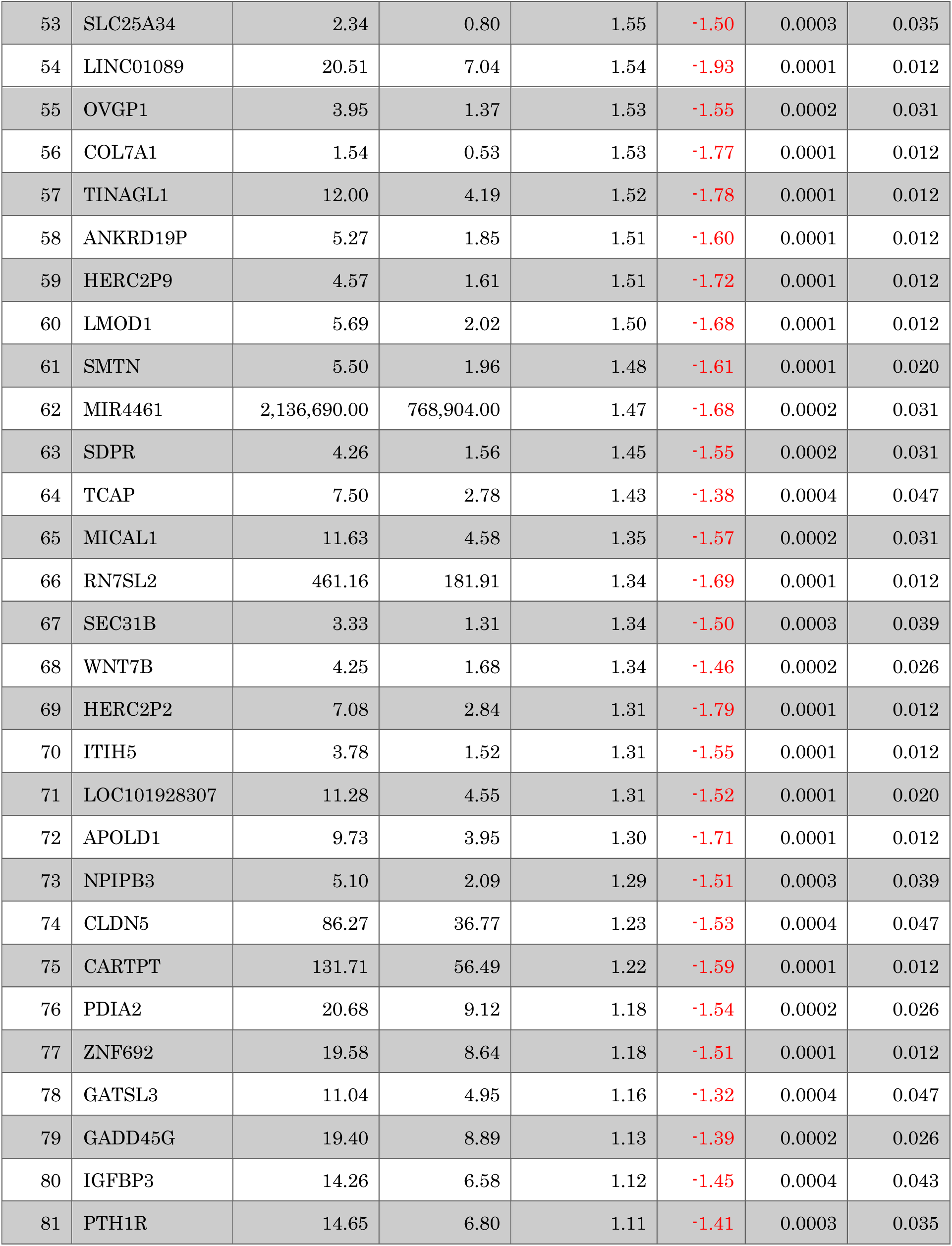
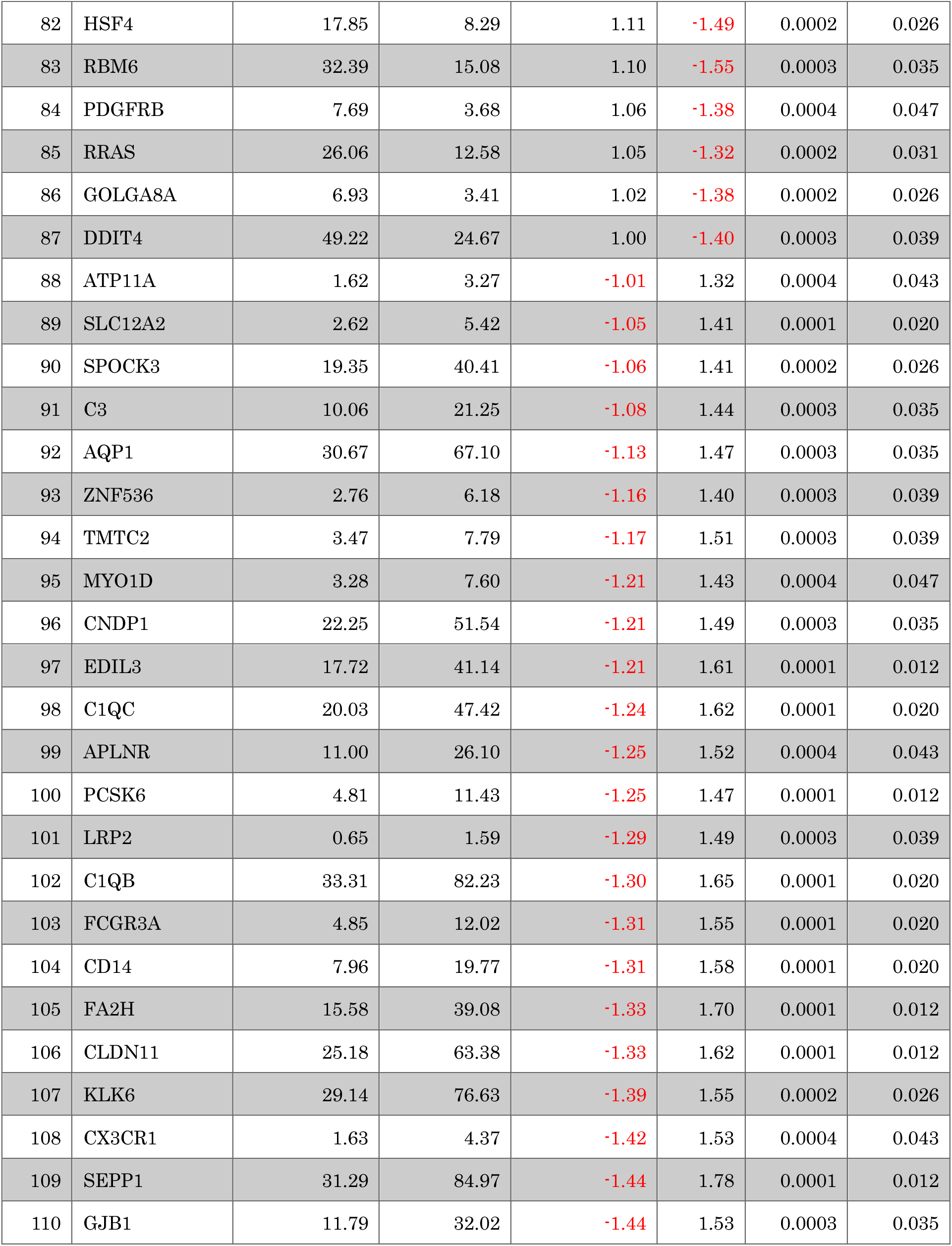
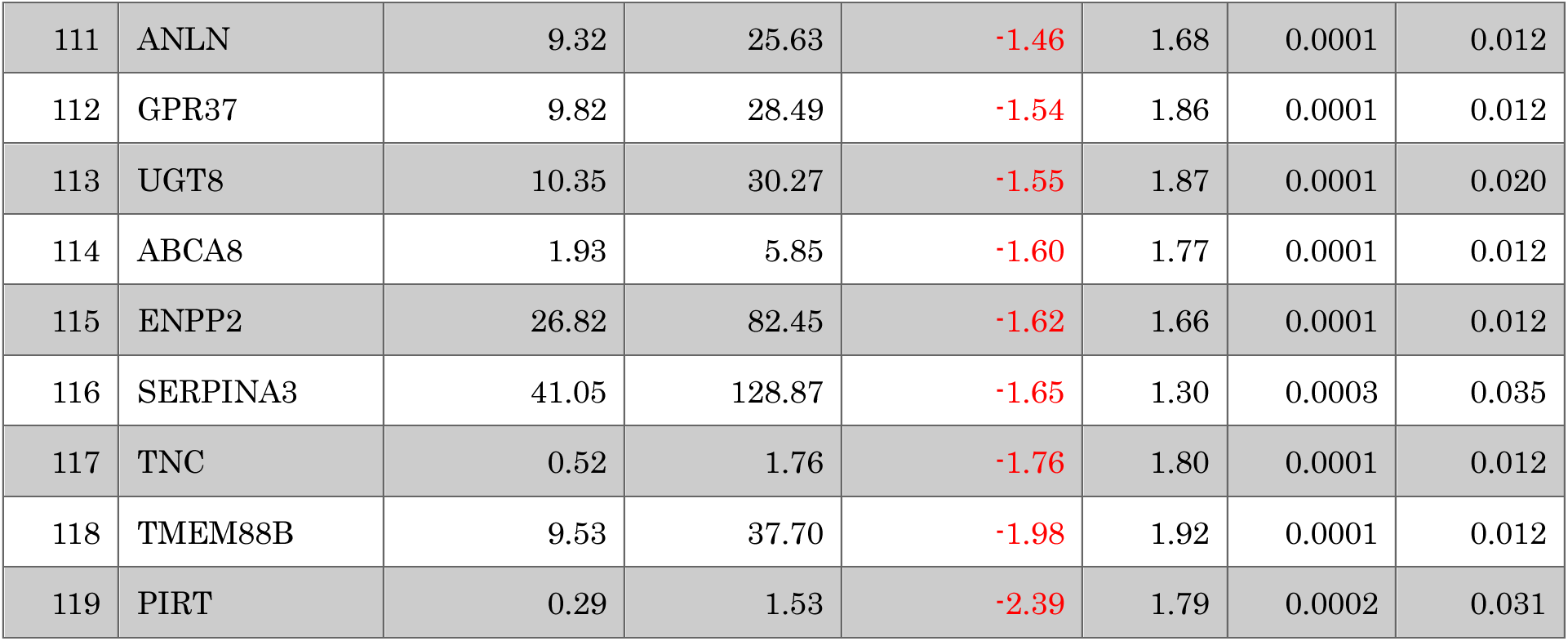
Differentially expressed genes in schizophrenia cases and controls. All of the DEGs were determined on the basis of statistical significance with FDR < 0.05 using Cuffdiff software. Cuffdiff uses a negative binomial distribution with generalized linear models to determine significance.

## Notes

### Competing Interest Statement

The authors have declared no competing interest.

